# Nutrient supplementation experiments with saltern microbial communities implicate utilization of DNA as a source of phosphorus

**DOI:** 10.1101/2020.07.20.212662

**Authors:** Zhengshuang Hua, Matthew Ouellette, Andrea M. Makkay, R. Thane Papke, Olga Zhaxybayeva

**Affiliations:** Department of Biological Sciences, Dartmouth College, Hanover, NH, USA; Department of Molecular and Cell Biology, University of Connecticut, Storrs, CT, USA; Department of Computer Science, Dartmouth College, Hanover, NH, USA

**Keywords:** extracellular DNA, dissolved DNA, hypersaline, haloarchaea, Halobacteria, DNA uptake, community diversity

## Abstract

All environments including hypersaline ones harbor measurable concentrations of dissolved extracellular DNA (eDNA) that can be utilized by microbes as a nutrient. However, it remains poorly understood which eDNA components are used, and who in a community utilizes it. For this study, we incubated a saltern microbial community with combinations of carbon, nitrogen, phosphorus, and DNA, and tracked the community response in each microcosm treatment via 16S rRNA and *rpoB* gene sequencing. We show that microbial communities used DNA only as a phosphorus source, and provision of other sources of carbon and nitrogen was needed to exhibit a substantial growth. The taxonomic composition of eDNA in the water column changed with the availability of inorganic phosphorus or supplied DNA, hinting at preferential uptake of eDNA from specific organismal sources. Especially favored for growth was eDNA from the most abundant taxa, suggesting some haloarchaea prefer eDNA from closely related taxa. Additionally, microcosms’ composition shifted substantially depending on the provided nutrient combinations. These shifts allowed us to predict supplemented nutrients from microbial composition with high accuracy, suggesting that nutrient availability in an environment could be assessed from a taxonomic survey of its microbial community.

## Introduction

Dissolved extracellular DNA (eDNA) in the environment plays important roles in global cycles of carbon (C), nitrogen (N) and phosphorus (P). It is found at measurable concentrations in every analyzed habitat [1–4] and is estimated to amount to gigatons globally [5]. Presence of eDNA in environments is predominantly influenced by release of DNA via microbial cell lysis from viral infections [6, 7], but also results from DNA freed up from spontaneous and programmed cell death, autolysis, and direct DNA secretion [8–11]. Subsequently, eDNA becomes available for resident organisms to utilize, and in some environments eDNA turnover rates can be as short as a few hours [12]. However, seasonal variability and type of habitat can increase the residence time of eDNA to months and years [13, 14], despite the presence and high activity of secreted extracellular nucleases [3]. Adsorption to minerals and particles can make eDNA unavailable for microorganisms [15], but factors like nutrient obtainability, and the presence of salt could also contribute to the stability, availability and utilization of the available eDNA [13, 16].

Many microorganisms secrete enzymes that degrade eDNA extracellularly [17]. The subsequent uptake of the DNA components could be used to support growth by being used as a C, N or P source [18, 19]. Many archaea and bacteria can also take up double-stranded eDNA as a high molecular weight molecule via a process of natural transformation, and use it either to repair damaged chromosomes or to increase genetic variation [15, 20]. However, since natural transformation has low efficiency rates and only one DNA strand is typically used in recombination, most of the eDNA transported into the cell may be consumed for other purposes, such as an energy source or as input building blocks (e.g., nucleotides) in metabolic pathways [21–25].

Hypersaline environments are reported to have the highest concentrations of eDNA [3]. These habitats have little to no primary production near the saturation level of NaCl, yet they support a high density of cells, usually dominated by heterotrophic, aerobic, obligate halophilic archaea, informally known as Haloarchaea [26–28]. A nutritionally competent model haloarchaeon *Haloferax volcanii* can grow on eDNA, primarily as a P source [29], suggesting that in hypersaline habitats eDNA plays an important role in the phosphorus cycle and also possibly in carbon and nitrogen cycles. Notably, *Hfx. volcanii* discriminates different eDNA sources for growth, as cultures did not grow on purified *E. coli* or herring sperm DNA, but grew on *Hfx. volcanii* (i.e., its own) or unmethylated *E. coli* DNA [23]. Although some bacteria have demonstrated biased uptake of DNA for natural transformation purposes [30–33], the above observations for *Hfx. volcanii* raise the possibility that many other microorganisms in a haloarchaeal community can grow only on specific eDNA, perhaps based on its methylation status. This remains a largely unexplored phenomenon. To better understand how eDNA influences growth of microbial communities in hypersaline environments, and to investigate eDNA utilization biases associated with organismal source of eDNA by different haloarchaea and hypersaline-adapted bacteria, we conducted microcosm experiments on natural near-saturated hypersaline waters collected from the Isla Cristina solar saltern in southern Spain that were amended by sources of C, N, inorganic phosphorus (Pi) and DNA. We show that eDNA, which was either available in the water column or provided as a supplement, was utilized by the microbial community as a source of phosphorus. Via sequencing of *rpoB* and 16S rRNA genes from DNA collected from both the cells and water column before and after the experiments, we observe that composition of both microbes in the community and eDNA in the water column changes depending on nutrients, and infer that at least some of these shifts are due to the ability to use eDNA and to prefer eDNA of specific taxonomic origin or taxonomic relatedness. We also found that the shifts in taxonomic composition of a saltern microbial community and eDNA in the water column are substantial enough to be predictive of the provided nutrients.

## Materials and Methods

### Nutrient supplementation of samples, microcosms’ incubation, DNA purification, rpoB and 16S rRNA genes amplification

Isla Cristina water samples were aliquoted into 50mL conical tubes, with each tube receiving 30mL of water sample. These samples were supplemented with various combinations of C (0.5% w/v glucose), N (5 mM NH_4_Cl), and P_i_ (1 mM KH_2_PO_4_) sources, as well as with high molecular weight, undigested genomic DNA (~6ng/μL of either *Hfx. volcanii* DS2 or *E. coli dam*^*−*^/*dcm*^*−*^) (the listed concentration values are final concentrations). Each combination of nutrient supplementation was performed in duplicate. The microcosms were incubated at 42°C and once they reached stationary phase, they were centrifuged and filtered. DNA was extracted, and *rpoB* and 16S rRNA genes were PCR-amplified. For details see **Supplementary Methods.**

### Quantification of nutrients in Isla Cristina water samples

The Isla Cristina water samples before and after experiments (including duplicates) were analyzed for total organic carbon (TOC), total dissolved nitrogen (TN), total phosphorus (TP) and inorganic phosphorus (P_i_). For each microcosm, 5 mL was aliquoted into 50 mL conical tubes, which were then diluted 1:10 with deionized water. The organic phosphorus in each sample (P_o_) was calculated as the difference between TP and P_i_. See **Supplementary Methods** for details.

### DNA sequencing, sequence quality control, pair merging, clustering of sequences into the operational taxonomic units and assignment of taxonomic affiliations

The purified amplicon products were used to construct 69 *rpoB-*based and 28 16S-rRNA-based libraries. The libraries were sequenced using Illumina MiSeq technology, collectively producing 3 567 797 and 2 277 899 paired-end raw reads for *rpoB* and 16S-rRNA genes, respectively. The raw reads were trimmed from poor quality regions and pairs were merged as described in **Supplementary Methods**. This procedure resulted in the 1 174 289 *rpoB* and 1 320 673 16S *rRNA* merged pairs, which were clustered into operational taxonomic units (OTUs) using QIIME v1.9.1 [34] and assigned taxonomic affiliations using the Ribosomal Database Project classifier v2.2 [35], as detailed in **Supplementary Methods**. Since *E. coli* and *Hfx. volcanii* are not indigenous members of the initial microbial community, the DNA assigned to these taxa were excluded from all further analyses.

### Quantification of similarities and differences in taxonomic composition across and within samples

For each sample, relative abundance of its OTUs, overall taxonomic composition and the OTU diversity were calculated as described in **Supplementary Methods**. Additionally, the samples were clustered via <u>P</u>rincipal <u>Co</u>ordinates <u>A</u>nalysis (PCoA), using beta-diversity distances calculated from Bray-Curtis and Jaccard dissimilarity values. The clusters were tested for similarity and differences in the OUT composition (see **Supplementary Methods** for details).

### Examination of the impact of provided nutrients on community composition using linear regression

Linear regression modeling was conducted with provided nutrients (C, N, P_i_ and P_o_) as explanatory variables and number of OTUs in each sample as dependent variables as described in **Supplementary Methods**.

### Quantification of changes in relative abundances of the abundant haloarchaeal OTUs (ahOTUs) in response to provided phosphorus

Changes in ahOTU abundance in microcosms and eDNA pools in response to addition of DNA and P_i_ were visualized as heatmaps mapped to a cladogram of relationships among ahOTUs (see **Supplementary Methods** for details). Additionally, changes in relative abundance of an ahOTU in microcosms and eDNA pools between two experimental treatments were summarized using two metrics, *D* = (*ICW*_*a*_ − *ICW*_*b*_) − (*ICC*_*a*_ − *ICC*_*b*_) and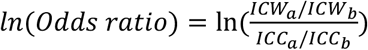, where *a* and *b* denote two compared treatments and *ICC*_*i*_ and *ICW*_*i*_ represent relative abundances of an ahOTU in communities and eDNA pools of a treatment *i*, respectively.

### Prediction of provided nutrients from taxonomic composition of a sample

Using the relative abundance of ahOTUs in a sample and categorization based on combination of nutrients provided in the experiments, a random forest classifier [36] was trained and cross validated as described in **Supplementary Methods**. The performance of classifier trained on the real data was compared to that of a classifier trained on the re-shuffled samples.

### Data availability

Raw sequencing data is available in GenBank under BioProject ID PRJNA646965. **Supplementary Data**, which includes a script used for raw data quality control, merged pairs, database used for *rpoB*-based OTU taxonomy assignment, tables of OTU composition per sample, and nucleotide sequences chosen to represent each OTU, are available in FigShare repository at https://doi.org/10.6084/m9.figshare.12609068.

## Results

### Isla Cristina’s microbial communities grow on eDNA as a phosphorus source

Supplementation of microbial communities from the Isla Cristina saltern (Huelva, Spain) with various combinations of C, N, P_i_, and DNA from either *Hfx. volcanii* DS2 (denoted as H) or *E. coli dam*^*−*^/*dcm*^*−*^ (denoted as E) revealed that unless both C and N were provided, no notable growth was observed (**Fig. 1a** and **1b**). Supplementation with both C and N (+C+N treatment) yielded an average net OD_600_ increase of 0.414 (**Fig. 1a**) and 0.506 (**Fig. 1c**) in the two sets of conducted experiments. Providing P_i_ in addition to C and N (+C+N+P_i_ treatment) resulted in the largest growth (**Fig. 1a** and **Fig. 1c**). Interestingly, samples that were supplemented with DNA instead of P_i_ (+C+N+H or +C+N+E treatments) also exhibited a comparable amount of growth, although net OD_600_ increase was ~12.9% lower than in the +C+N+P_i_ treatment (One-way ANOVA test; P-value 0.0036; **Fig. 1c**). Nevertheless, the net OD_600_ increases in the +C+N+H and +C+N+E treatments were 50% and 54% higher, respectively, than in the +C+N treatment (One-way ANOVA test; P-values 0.0023 and 0.0018, respectively; **Fig. 1c**). This suggests that besides P_i_, the saltern community can utilize eDNA as a source of phosphorus.

**Figure 1.**
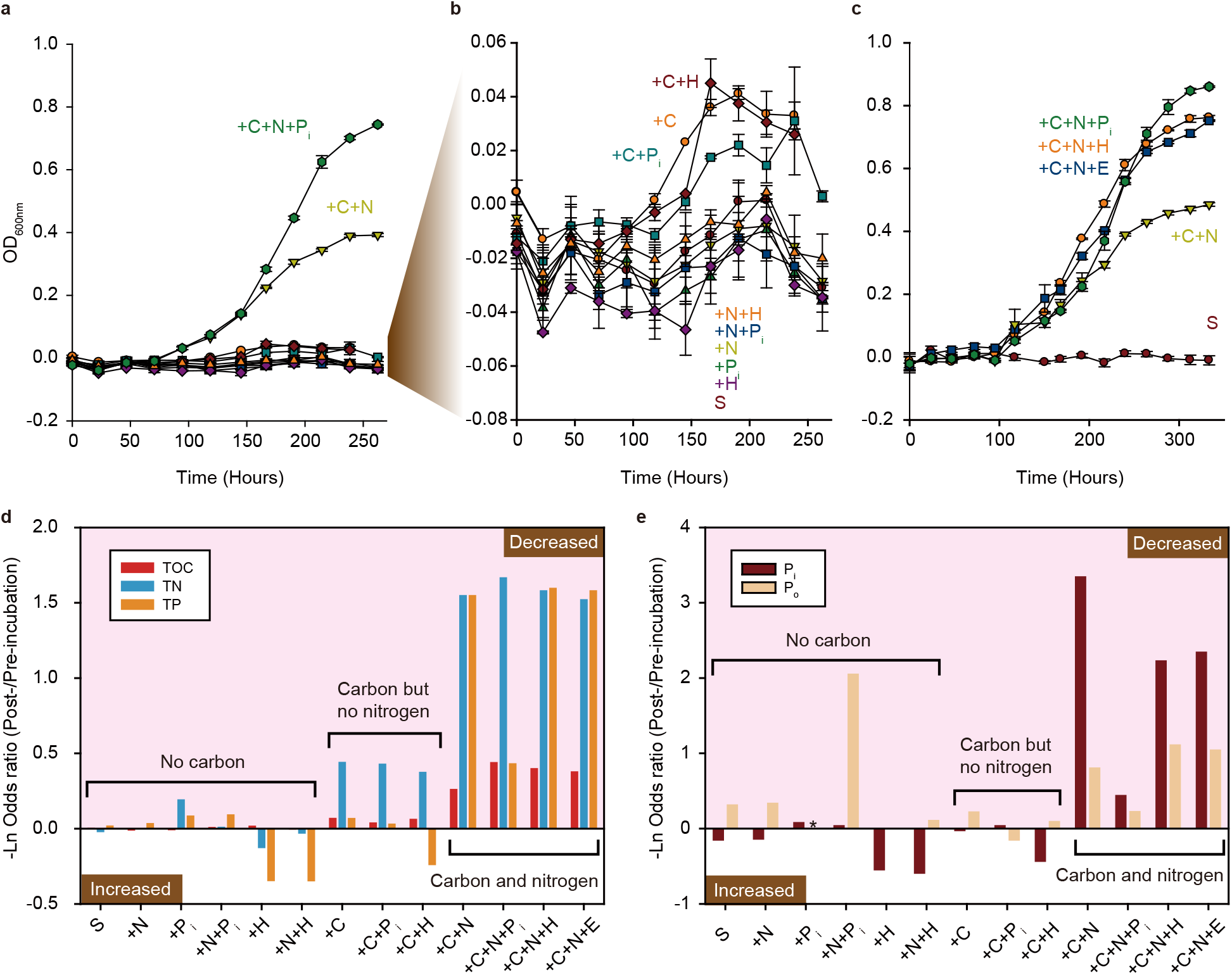
Microbial growth and associated changes in available nutrients in the experimental microcosms. **(a-c)** Growth curves under experimental treatments. Each line represents a separate experimental treatment. For each treatment, the added nutrients are denoted as “+” followed by a nutrient symbol of carbon (C), nitrogen (N), phosphorus (P_i_), *Hfx. volcanii* DS2 DNA (H) and *E. coli dam*^*−*^/*dcm*^*−*^ DNA (E). Starvation growth curve (S) serves as a control. Error bars represent the standard error of the mean. **Panel b** shows growth curves of the slow-growing microcosms from **panel a** plotted on a different scale. **Panels a** and **c** represent two independent sets of experiments. **(d, e)** Change in carbon, nitrogen and phosphorus after the microbial growth has stopped. The plotted changes are mean values from two experimental replicates. The experimental treatments (X axis) are grouped into three categories based on whether carbon and/or nitrogen were provided. Nutrient abbreviations: TOC, total organic carbon; TN, total nitrogen; TP, total phosphorus; P_i_, inorganic phosphorus. Organic phosphorus (P_o_) was computed as by subtracting P_i_ from TP. For “+P_i_” treatment, the calculated value of P_o_ is replaced with an asterisk, due to errors associated with measurements of TP and P_i_ for that treatment. Actual values for the measured nutrient concentrations and the associated calculations are shown in **Supplementary Tables S1** and **S2**.

The observed patterns of microbial community growth under different treatments are consistent with the consumed amounts of C, N, and P_i_, as measured by comparing the total organic carbon (TOC), total nitrogen (TN), and total phosphorus (TP) in the filtered untreated water samples and in the treated filtered water samples after the communities reached the stationary phase. The pre-incubation communities contained small quantities of TOC, TN, and TP (**Supplementary Table S1**). In treatments where communities exhibited no notable growth (i.e., C was not provided), there were also no notable changes in TOC, TP and TN (**Fig. 1d**). In the microcosms with some growth (C was provided but N was not), a 31-36% decrease in TN was observed (**Fig. 1d** and **Supplementary Table S1**), suggesting that growth was sustained until naturally occurring utilizable nitrogen resources were depleted. The fastest growing microcosms, i.e. in which both C and N were supplied, exhibited the largest decreases in TN, TP and TOC (**Fig. 1d**).

To further investigate the effectiveness of DNA as a source of phosphorus, we examined the contributions of inorganic (P_i_) and organic (P_o_) phosphorus to the TP in the microcosms after they reached stationary phase (**Fig. 1e** and **Supplementary Table S2**). We found that in the microcosms supplemented with C and N but not provided with P_i_ (i.e., +C+N, +C+N+H or +C+N+E treatments), the remaining TP was dominated by P_o_ (~91%, ~70% and ~74%, respectively). Since the microcosms under the +C+N, +C+N+H or +C+N+E treatments exhibited substantial growth, we conjecture that while DNA (i.e., P_o_) was used by the communities as a source of phosphorus, P_i_ is generally preferred when available.

### The taxonomic composition of the microbial communities and eDNA pool varies across treatments

To better understand how the different nutrient availability affects both organismal composition of a community and organismal sources of DNA in the eDNA pool, we sequenced 16S rRNA and *rpoB* genes from the DNA extracted from the <u>I</u>sla <u>C</u>ristina living <u>C</u>ells (ICC) and the <u>I</u>sla <u>C</u>ristina <u>W</u>ater column (ICW) before and after the above-described microcosm experiments.

Analysis of the 16S rRNA OTUs revealed that, as expected for a saltern environment, the pre-incubation community (ICC sample “X” on **Fig. 2a**) is numerically dominated by archaea from class Halobacteria (relative abundance of 92.9%) and by bacteria from the genus *Salinibacter* (relative abundance of 6.6%). The remaining 0.5% of OTUs belong to either other archaeal classes or other bacterial genera (see **Supplementary Data**). Halobacteria and *Salinibacter* also dominate all post-treatment communities, with their relative abundances in 92.3-99.9% and 0.02-7.2% range, respectively (**Fig. 2a**).

**Figure 2.**
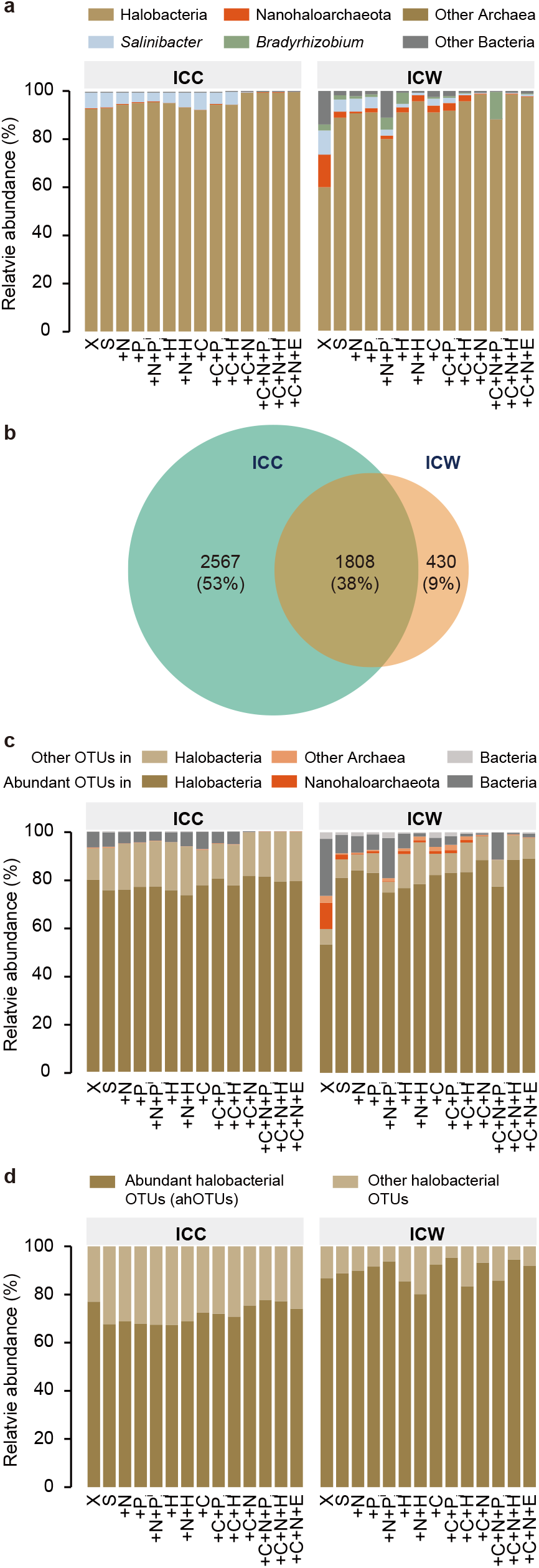
OTU composition of the microbial communities (ICC) and of eDNA in the associated water columns (ICW). **(a)** Relative abundance of 16S rRNA-based OTUs in the pre-incubation community (X), after starvation (S), and after nutrient-addition experiments. For experimental treatment abbreviations see **Figure 1** legend. Only a few selected taxonomic groups are highlighted, while other OTUs are pooled into “Other Archaea” and “Other Bacteria” categories. **(b)** Overlap of 16S rRNA-based OTUs in ICC and ICW across all samples combined. **(c)** Relative abundances of 16S rRNA-based OTUs that constitute ≥ 1% of at least one sample (designated as “abundant OTUs”) in comparison to the OTUs with < 1% abundance (denoted as “Other OTUs”). **(d)** Relative abundances of *rpoB*-based OTUs from class Halobacteria that constitute ≥ 1% of at least one sample (designated as “ahOTUs”) in comparison to the Halobacterial OTUs with < 1% abundance (denoted as “Other Halobacterial OTUs”).

The DNA in water column is also dominated by eDNA from Halobacteria and *Salinibacter* OTUs (**Fig. 2a**). However, DNA originating from bacteria on average represents a higher fraction in the eDNA pools than in the microbial communities, with the largest fraction (27%) found in the pre-incubation water column (ICW sample “X” on **Fig. 2a)**. Additionally, some eDNA comes from OTUs that were either not observed in any of the ICC samples (**Fig. 2b**) or were found in the ICC samples in low abundances. Examples include OTUs from bacterial genus *Bradyrhizobium* and archaeal subphylum Nanohaloarchaeota (see **Supplementary Results** and **Supplementary Data** for details). These eDNA fragments may have originated from living microorganisms elsewhere and were transported to the saltern environment. Alternatively, they may represent resistant-to-consumption eDNA that has slowly accumulated from the rare members in the pre-incubation community (see **Supplementary Results** for specific examples.)

Overall, OTUs present at >1% abundance constitute the greatest majority of OTUs in every sample (**Fig. 2c**). Unfortunately, many haloarchaea are known to carry multiple non-identical copies of 16S rRNA gene [37–39], with some copies exhibiting within-genome identity as low as 91.3% (**Supplementary Table S5**). Since this intragenomic heterogeneity is likely to affect accuracy of the 16S rRNA-based relative abundance estimates, we also amplified the *rpoB* gene, which is known to exist in haloarchaeal genomes in a single copy, and was shown to resolve genera assignments better than the 16S rRNA gene [40–42]. Because the 16S rRNA analysis indicates that the microbial communities are dominated by Halobacteria and do not contain any OTUs from other archaeal classes at > 1% abundance, we amplified the *rpoB* gene using the Halobacteria-specific primers. Of the 5 905 *rpoB*-based OTUs detected across all samples, 5 851 (99.1%) are assigned to class Halobacteria, demonstrating that the primers designed to amplify this class of Archaea have high specificity. Relative abundances of Halobacterial genera observed in the *rpoB*-based and 16S rRNA-based analyses are comparable, with few discrepancies detailed in **Supplementary Results**.

Thirty-four of the *rpoB*-based OTUs have > 1% relative abundance in at least one ICC sample. Thirty two of the 34 OTUs maintain > 1% relative abundance in at least one ICW sample. Additional 29 halobacterial OTUs have > 1% relative abundance in at least one ICW sample, and found in low abundance in at least one ICC sample. For the remainder of the analyses we designated these 34 + 29 = 63 OTUs as *<u>a</u>bundant <u>h</u>alobacterial OTUs* (ahOTUs). Notably, an individual sample contains only a fraction of the 63 ahOTUs: 16 ± 1.8 of ahOTUs per ICC sample and 18 ± 2.8 ahOTUs per ICW sample. Because in each sample the cumulative relative abundance of ahOTUs is 67-78% and 80-95% for ICC and ICW samples, respectively (**Fig. 2d**), there are non-negligible differences in the ahOTU composition and their relative abundances across samples. Below, we examine in detail how these differences relate to nutrient availability.

### Carbon, DNA and inorganic phosphorus alter taxonomic composition of microbial communities and eDNA pools

Based on the relative abundances of both the *rpoB*-based and the 16S rRNA-based OTUs, the samples aggregated into three distinct clusters **(Fig. 3** and **Supplementary Fig. S1a)**. Differences in OTU composition of the three clusters are significant (PERMANOVA: R = 0.74, P = 0.001 [*rpoB*] and R = 0.72, P = 0.001 [16S rRNA]). Cluster 1 consists of the cellular communities’ samples that come from the pre-treatment and all microcosms that either exhibited no notable or some growth (**Fig. 1b**), hereafter referred as “slow-growing communities”. Cluster 2 consists of the water column samples that correspond to the cellular communities of Cluster 1. Cluster 3 consists of the cellular communities’ and water column samples from all experiments that produced fast-growing microcosms, that is, the microcosms that were supplemented simultaneously with C and N. Such clear taxonomic separation of fast- and slow-growing microcosms suggests substantial shifts in community composition in response to nutrient supplementation. Clusters 1, 2 and 3 also emerge even if OTU abundance is ignored and only OTU richness is considered (**Supplementary Fig. S2**), suggesting that the observed clustering is not driven solely by a few abundant community members, but also influenced by rare microorganisms and their DNA in the water column.

**Figure 3.**
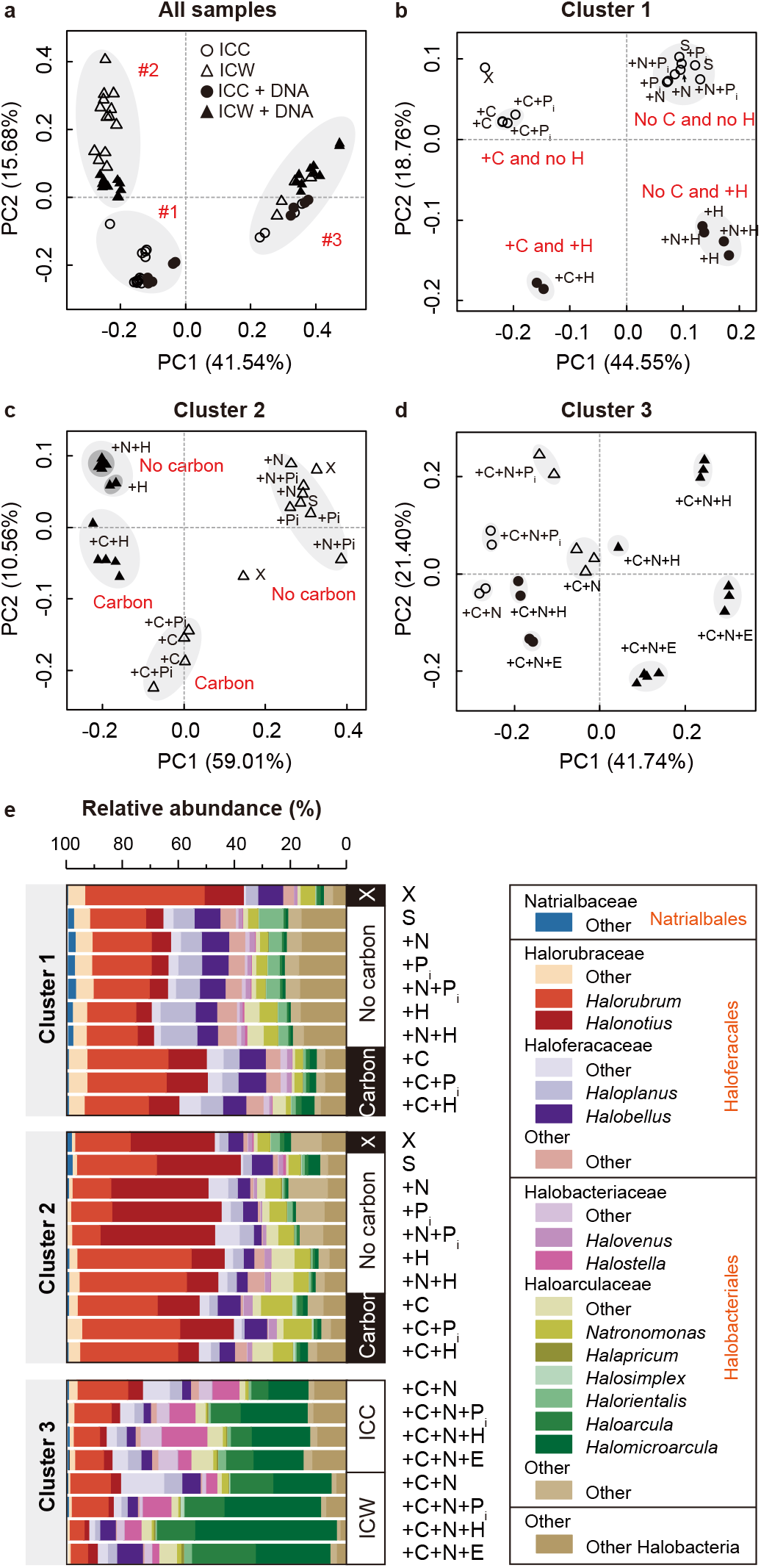
Comparison of the *rpoB*-based OTU compositions of the analyzed samples. (**a**) Principal Coordinate Analysis (PCoA) of all samples using the pairwise Bray-Curtis dissimilarity as a distance metric. The distances were calculated using all OTUs in a sample. Circles denote microbial community samples, while triangles refer to the water column samples. Within these, samples for treatments with added DNA are shown using filled symbols, while all remaining samples are shown using open symbols. For the PCoA analyses carried out using 16S rRNA-based OTUs, see **Supplementary Figure S2**. (**b**, **c** and **d**) PCoA of the samples within each of the three clusters. Same distance measure and same symbol notations as in panel **a**. (**e**) Taxonomic composition of samples within each of the three clusters. See **Supplementary Table S4** for the actual relative abundance values.

Presence and abundance of specific bacterial and archaeal OTUs, and especially those with > 1% abundance, are behind the observed shifts in taxonomic composition among the three clusters (**Fig. 2**). Among Halobacteria, the relative abundances of 48 of the 63 ahOTUs differ significantly among the three clusters (the Mann-Whitney *U* test; see **Supplementary Table S6** for P values; **Fig. 4a**). Cluster 1 samples are dominated mostly by Haloferacaceae ahOTUs (**Fig. 4b** and **4c**). In contrast, eDNA from Halorubraceae ahOTUs (**Fig. 4b** and **4d**) dominate Cluster 2 samples. Finally, in Cluster 3 samples, Haloferacaceae and Halorubraceae ahOTUs become much less abundant and are replaced by Haloarculaceae and Halobacteriaceae ahOTUs (**Fig. 4b** and **4e**).

**Figure 4.**
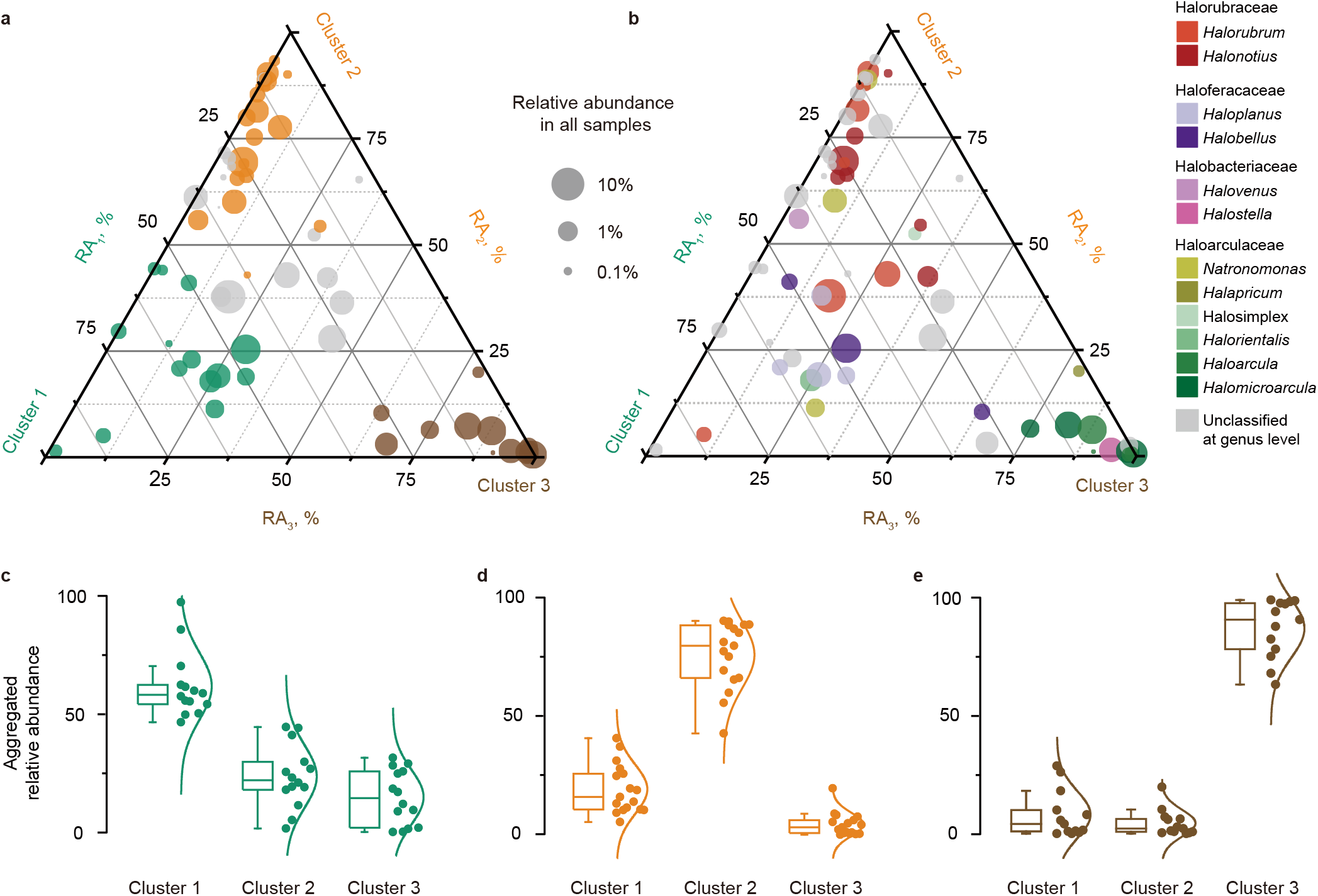
Relative abundance of ahOTUs across three clusters. (**a** and **b**) Relative abundance of ahOTUs in three clusters summarized in a ternary plot. Each vertex represents one of the three clusters. Each circle represents an ahOTU. The position of a circle is determined by relative abundance (RA) of the ahOTU in three clusters, and the circle size is proportional to the ahOTU’s average relative abundance across all 69 samples. In **panel a**, ahOTUs with significantly higher abundance in Cluster 1, 2 or 3 than in the other two clusters are colored in green, orange and brown, respectively, while OTUs without significant difference in abundance are shown in grey. In **panel b**, the circles are colored according to the taxonomic assignment of the ahOTUs. (**c, d** and **e**) Aggregated relative abundances (aRA) of ahOTUs with significantly higher abundance in one cluster across each of the three clusters. Each point represents an ahOTU. For each OTU its aRA in a cluster *i* (*i = 1,2,3*) is defined as 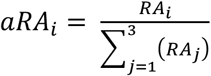, where *RA*_*i*_ is the relative abundance of the ahOTU in the cluster *i*. The distribution of aRAs within a cluster is summarized by a probability density function and a box-and-whisker plot, with whiskers extending to 1.5 of the interquartile range.

Within the three clusters, there are visibly distinct sub-clustering of samples that imply more nuanced differences in taxonomic composition of the samples. For the Cluster 1 communities and their water columns (Cluster 2), the pre-treatment sample, samples without notable growth (no C added) and some growth (C added) form separate clusters, with further subdivision due to provision of *Hfx. volcanii* DNA (PERMANOVA: R = 0.75 and 0.84, P = 0.001 and 0.001 [*rpoB*] and R = 0.74 and 0.81, P = 0.008 and 0.007 [16S rRNA]) (**Fig. 3b-c** and **Supplementary Fig. S1b-c**). Within fast-growing microcosms (Cluster 3), sub-clustering is less pronounced but reflects availability and type of P sources provided in treatments (**Fig. 3d** and **Supplementary Fig. S1d**). Taxonomically, the observed groups are due to differences in relative abundance of Halorubraceae, Haloarculaceae and Haloferacaceae OTUs for slow-growing communities and of Haloferacaceae, Halorubraceae, Haloarculaceae and Halobacteriaceae OTUs in fast-growing communities (**Fig. 3e**).

Taken together, supplementation of nutrients significantly influences composition of both the microbial community and the eDNA pool. Linear regression modeling with nutrients as categorical variables revealed that provision of C is the primary driver of microbial community composition shifts (R_adj_^2^ = 0.45; P = 0.002; *t* value: 4.5), suggesting unequal abilities of various taxa to compete for added glucose. Provision of N alone does not affect community composition, suggesting that community members have equal abilities in its assimilation. Provision of P_i_ and DNA significantly alters the composition of eDNA in the water column (R_adj_^2^ = 0.327; P = 0.001; *t* value: 3.6), raising a possibility that there are organismal preferences for phosphate and specific “taxonomic affiliation” of DNA as a P source.

### Shifts in eDNA OTU composition depending on available sources of phosphorus suggest preferential eDNA uptake

It is surprising that the DNA content of the initial community and the slow-growing microcosms (Cluster 1) are quite different from the eDNA pools of their water columns (Cluster 2). This distinction, albeit less apparent, also exists in the fast-growing microcosms (Cluster 3; **Fig. 3d**). Assuming that all cells in a microcosm have equal probability of birth and death, it would be expected to see similar OUT compositions of a microbial community and of the eDNA from the corresponding water column. Interestingly, in the slow-growing communities, provision of *Hfx. volcanii* DNA (+H and +C+H treatments) significantly increases the eDNA OTU richness (the Mann-Whitney *U* test, P = 1.6×10^−5^; **Supplementary Fig. S3**). In the fast-growing communities, the increase in eDNA OTU richness, albeit not significant, is caused by the addition of P_i_ (+C+N+P_i_ treatment; **Supplementary Fig. S3**). Therefore, we hypothesize that the observed differences in OTU compositions of cellular communities and their water columns are due to biased eDNA uptake from certain taxonomic groups. Furthermore, we propose that the availability of *Hfx. volcanii* DNA and inorganic phosphorus in slow- and fast-growing communities, respectively, results in the reduced uptake of available eDNA as a P source, leading to eDNA accumulation in the environment and causing the observed changes in eDNA OTU composition. And in reverse, the observed limited OTU diversity when P sources are scarce may be due to depletion of eDNA from specific taxa through taxonomically-biased eDNA uptake.

### Preferences for eDNA of specific taxonomic origins by the saltern microbial community

Complexity and variability of microcosms across experimental treatments (**Fig. 3e**), as well as impacts of sequencing depth and sequencing biases, make it difficult to establish exact eDNA preferences of specific OTUs from our data. However, by comparing relative abundances of specific OTUs in microcosms with and without additional P sources, we can indirectly infer some taxonomic biases of eDNA uptake in this saltern microbial community as a whole. Specifically, we will continue to assume that all cells in a microcosm have equal probability of birth and death. As a result, OTUs that do not substantially change their relative abundance in treatments with and without an extra P source are not expected to show substantial changes in the relative abundance of their eDNA in the water column. Deviations from these patterns would suggest an eDNA uptake bias, and we developed and applied two metrics to quantify such differences (see **Materials and Methods** for details).

From comparisons of slow-growing microcosms with and without added *Hfx. volcanii* DNA, we found that when *Hfx. volcanii* DNA is not provided, eDNA from five Halorubraceae and Haloarculaceae ahOTUs is depleted (see highlighted taxa in **Supplementary Fig. S4** and **Supplementary Table S7**) and eDNA of 305 additional OTUs become undetectable in the water columns (**Supplementary Table S8**). Of these 305 OTUs, 98 (32%) belong to Halorubraceae and Haloarculaceae families. These findings suggest preferential consumption of Halorubraceae and Haloarculaceae eDNA in slow-growing microcosms. In fast-growing microcosms, eDNA from Haloarculaceae, Halorubraceae and Halobacteriaceae ahOTUs are depleted in water columns without provided P source (either *Hfx. volcanii* DNA, *E. coli* DNA or P_i_; see highlighted taxa in **Supplementary Fig. S5** and **Supplementary Table S9**). In comparisons between +C+N and +C+N+P_i_ treatments, eDNA from 158 OTUs become depleted in the water column without P_i_, and 62 of the 158 OTUs (39%) belong to Haloarculaceae, Halorubraceae and Halobacteriaceae families (**Supplementary Table S10**). These observations suggest preferential consumption of eDNA from these three families in fast-growing microcosms. Interestingly, the most abundant of the depleted eDNA belongs to ahOTUs from families that represent a substantial fraction of the microcosms (Halorubraceae for the slow-growing and Haloarculaceae for the fast-growing communities; **Fig. 3e**), indicating that eDNA most closely related to the most abundant OTUs in the community is preferentially consumed. Intriguingly, several ahOTUs become depleted in the +C+N+P_i_ water columns when compared to the other three treatments with fast growth (+C+N, +C+N+H, and +C+N+E) (**Supplementary Fig. S5** and **Supplementary Table S9**), suggesting that some eDNA may be more preferred by some organisms than inorganic phosphorus.

We also hypothesize that the saltern community can also utilize bacterial eDNA, which can be exemplified by Chitinophagaceae DNA (**Supplementary Fig. S6**). Chitinophagaceae OTUs are not detected in the pre-incubation cellular community sample, but their DNA is abundant in the pre-incubation water column. Under all experimental treatments, the Chitinophagaceae OTUs continue to be undetected in the microbial communities, suggesting they are at best extremely rare community members. However, their DNA is either absent or found at very low abundance in the treatments’ water columns, suggesting that Chitinophagaceae eDNA in the initial, pre-incubation water column was consumed by the microcosms’ communities.

Provision of large amounts of *Hfx. volcanii* DNA allowed us to identify taxa that likely prefer to utilize *Hfx. volcanii* DNA as a P source (**Supplementary Table S7**). For example, the ahOTU *denovo3413*, which is assigned to *Haloarculaceae* family and has 91.4% of nucleotide identity to *Halapricum* sp. strain SY-39 (MF443862.1), has the average relative abundance of 0.08% in the slow-growing communities without provided *Hfx. volcanii* DNA and 0.05% in the corresponding water columns. However, when *Hfx. volcanii* DNA is supplied, the abundance of *denovo3413* increases significantly, to 4.6% in microbial communities and to 4.3% in the water column samples (the Mann-Whitney *U* test, P = 7.4×10^−5^ and 1.1×10^−5^, respectively).

The above inferences do rely on an assumption that all cells in a microcosm have equal probability of birth and death, which may not hold. Future work that establishes relative growth rates of individual OTUs in the community will allow more precise estimation of eDNA preferences.

### Provided nutrients can be predicted from taxonomic affiliations of microbial community members and of water column DNA

The shifts in community composition in response to availability of C, N, and P suggest that nutrients provided in the experiments may be predictable from the OTU composition associated with DNA in cells and dissolved in water columns. We designed a random forest classifier that takes ahOTU composition as an input and predicts if a sample falls into one of the five treatment groups: “No C was provided (no C)”, “C but no N were provided (+C)”, “Only C and N were provided (no P)”, “C, N and P_i_ were provided (+P_i_)”, or “C, N and DNA were provided (+H/+E)”. We found that a random forest classifier achieves 95.5% accuracy in a 20-fold cross-validation and 94.6% accuracy in out-of-bag (OOB) error estimate. These predictions have significantly higher accuracies than those obtained in the classifier trained on the randomly shuffled samples (Student’s *t* test, P < 2.2*10^−16^ for cross validation approach [**Fig. 5a**] and Kolmogorov-Smirnov test, P < 2.2*10^−16^ for OOB approach [**Fig. 5b**]). Predictions remain robust even for smaller training data sets (**Fig. 5c**). Overall, 63 ahOTUs can predict the available nutrients with the mean prediction accuracy of 62%. The predictions are strongly influenced by 24 ahOTUs where each one contributes ≥1% of the prediction accuracy (**Fig. 5d**) and their removal decreases the overall mean classifier accuracy by 49%. Consistent with this observation, 23 of the 24 ahOTUs have significantly different abundances in samples from the five groups (**Fig. 5e**; one-way ANOVA test, all 23 P values < 0.01).

**Figure 5.**
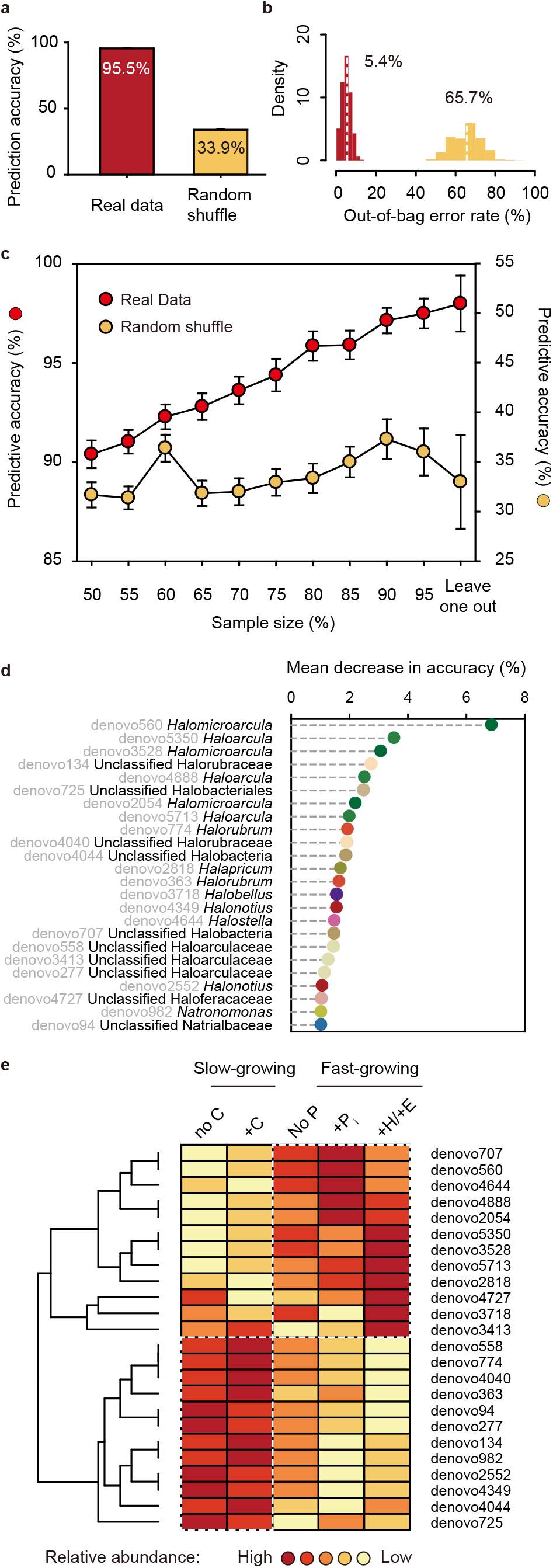
Performance of a random forest classifier in predicting available nutrients from the OTU composition of a microbial community. (**a**) Prediction accuracy in the 20-fold cross-validation of the classifier across 1000 training/validation datasets constructed from real and randomly reshuffled data. (**b**) Distribution of the out-of-bag (OOB) errors across 1000 training/validation datasets constructed from real and randomly reshuffled data. (**c**) Changes in prediction accuracy with the dataset size, which is shown as percent of the 69 samples used and as the leave-one-out strategy. (**d**) The decrease in prediction accuracy of 24 ahOTUs with at least 1% contribution to the predicted accuracy. The OTUs are colored according to notations shown in **Figure 3**. (**e**) The relative abundance of the top 24 ahOTUs in five categories of treatments. For each OTU, the relative abundance values were transformed into heatmap colors. The OTUs are clustered based on their relative abundance across five categories (as described in **Methods**). Abbreviations of five treatment categories: “+C”, C but no N were provided; “no C”, No C was provided; “+H/+E”, C, N and DNA was provided; “+P_i_”, C, N and P_i_ were provided; “No P”, only C and N were provided.

## Discussion

Our microcosm experiments with saltern-derived microbial communities revealed that carbon, nitrogen and phosphorus are limiting nutrients of the microcosms, and by inference the Isla Cristina crystallizer pond habitat, since their simultaneous provision resulted in the largest growth of cells. Having a readily accessible carbon source is most crucial for this microbial community, because no notable growth was observed without supplementation of carbon. Nitrogen, on the other hand, was more easily assimilated from the naturally available sources, as microcosms with no supplemented nitrogen exhibited growth until the environmental nitrogen was depleted. Interestingly, the communities supplied with carbon and nitrogen grew well even if not supplied with inorganic phosphorus, suggesting that they were able to utilize either provided DNA or naturally occurring eDNA as a source of phosphorus. However, eDNA (while likely present in all experimental microcosms) evidently could not serve as a source of carbon and nitrogen, for an unknown reason.

Nutrient usage of eDNA is not unique to the saltern microorganisms, as other microbes are known to use DNA as a nutrient by enzymatic cleavage of eDNA (either extracellularly or intracellularly) into deoxyribose, inorganic orthophosphate, and nucleobases [14, 17]. Yet, there is also a possibility that the observed growth in our microcosms was instead due to utilization of internal cellular DNA as a source of phosphorus. For example, previous experiments on a model haloarchaeon *Hfx. volcanii* demonstrated that in the absence of supplemented phosphate *Hfx. volcanii* was able to utilize phosphate stored in dozens of its chromosome and plasmids copies [23, 29]. Polyploidy was detected in all tested species and strains of Haloarchaea, and may be common in Haloarchaea [43–45] and Euryarchaeota [46]. Hence, many microorganisms populating the communities in our experiments could be polyploid and consequently could use the extra copies of their chromosomes as a P source.

Given the presumed utilization of eDNA as a P source, it is puzzling why so much of the available eDNA has remained unused. One possibility is that the community reached the stationary phase because it ran out of other nutrients first. For instance, nitrogen was used up in the fast-growing communities (**Supplementary Table S1**). Alternatively, the microorganisms may not have been able to utilize all available eDNA. For instance, aquatic bacteria are hypothesized to specialize on different sizes of DNA fragments [19], and naturally competent *Haemophilus influenzae* and *Neisseria gonorrhoeae* use species-specific DNA uptake recognition sequences and therefore would not import each other’s DNA [47–49]. DNA methylation patterns are also important for discriminating different sources of DNA during the uptake process [23, 50]. As a result, some eDNA might not be recognized as a P source. Consistent with these observations, the changes in eDNA composition of the water columns associated with our experimental microcosms showed that organismal origin of eDNA indeed matters for growth on eDNA, with some bacterial and archaeal eDNA being clearly preferred. Especially preferred is eDNA from the most dominant members of the microcosms, suggesting some haloarchaea prefer eDNA that is most closely related to their own. This preference may be based on DNA methylation patterns [23]. Future studies are needed to better understand the specific reasons behind the observed biases in eDNA utilization.

The observed patterns of preferential eDNA consumption and differential growth under various nutrient availability regimes are associated with substantial shifts in the taxonomic composition and overall diversity of underlying microcosm communities. While the “bottle effect” [51, 52] can never be completely ruled out for any laboratory microcosm experiments, lack of growth and absence of rapid changes in nutrient availability in the microcosms without provided nutrients, as well as similarity of the pre-incubation and slow-growing communities, suggest that the observed substantial changes to community composition occur in response to the addition of specific nutrient combinations rather than due to the bottle effect. Therefore, we conjecture that in salterns the microbial community assembly is determined by the available resources.

Such resource-based assembly also has been demonstrated in synthetic bacterioplankton communities [53], and therefore may represent a general principle for a microbial community assembly. The changes in a microbial community structure in response to environmental challenges are highly dynamic [54], and are likely due to a combination of differential abilities of lineages to survive under severe nutrient limitation [55, 56], to compete for assimilation of limited nutrients [57], to compete for uptake of abundant nutrients [58], to inhibit growth of their competitors [59], and to create mutualistic relationships via metabolic exchange and cross-feeding [54, 60]. Although relative importance of these factors in our saltern community remains to be established, differential abundance of Haloferaceae, Halorubraceae and Halobacteriaceae families in response to availability of carbon and phosphorus indicate that competition for the available nutrients plays a major role in shaping saltern communities. Moreover, after addition of limiting nutrients, the microcosms exhibited increased biomass production and reduction in OTU richness. Such loss of species diversity in response to increase in nutrient availability has been also demonstrated in bacterial biofilms [61], soil microbial communities [62], phytoplankton [63] and plants [64], and was interpreted as a support for the niche dimensionality theory [65]. The theory postulates that biodiversity is maintained under limiting resources because competition for them can be achieved via different trade-offs, which in turn creates multiple niches that can be occupied by similar but not identical species or lineages. Removing nutrient limitation eliminates trade-offs and leads to a decreased diversity. Yet, presence of several closely related and abundant OTUs from *Haloarcula* and *Halomicroarcula* genera in the fast-growing microcosms cannot be explained by competition under the niche dimensionality framework. Therefore, other processes, such as mutual exchange of metabolic by-products, may be at play to promote their coexistence [60]. Future metagenomic surveys of metabolic genes within these microcosms will help to test this hypothesis.

Changing microbial communities can in turn have a great impact on the environment they inhabit [66, 67]. By applying a machine learning algorithm to OTU abundance and nutrient treatments data from our microcosm experiments, we found that the nutrient availability could be predicted with high accuracy from the DNA composition of a microbial community and water column. Notably, to achieve the high accuracy it is sufficient to have data from only few abundant OTUs. Our findings suggest that nutrient availability in a natural environment (or its past and future changes) could be predicted from a taxonomic survey of a microbial community, if appropriate training data from laboratory experiments is available. This could become a useful tool for advancing our understanding of ecological processes happening in particular environments and for practical applications, like bioremediation projects or treatments of microbiome-impacted diseases.

## Supporting information

Supplementary Methods, Resulsts and Figs S1-S7

## Acknowledgements

We thank Drs. Antonio Ventosa, Paulina Corral, and Cristina Sánchez-Porro (University of Sevilla), who helped with saltern sampling. We also thank Dr. Jialiang Kuang (University of Oklahoma) for discussions about random forest predictions, and Dr. Shaopeng Li (East China Normal University) for discussions about community assembly and niche dimensionality. The work was supported through a NASA Exobiology award NNX15AM09G to R.T.P. and O.Z., and by the Simons Foundation Investigator in Mathematical Modeling of Living Systems award 327936 to O.Z.

## Author contributions

Z.S.H., O.Z., and R.T.P. conceived the study. M.O. and A.M. performed the microcosm experiments, DNA extraction and sequencing. Z.S.H. performed the metagenomic and statistical analyses. Z.S.H., O.Z., R.T.P. and M.O. wrote the manuscript. All authors discussed the results and commented on the manuscript.

