## Supplementary Methods, Resulsts and Figs S1-S7 for "Nutrient supplementation experiments with saltern microbial communities implicate utilization of DNA as a source of phosphorus"

**for**

**This file includes:**

- Supplementary Figures S1-S7.
- Supplementary Results.
- Supplementary Methods.

### Supplementary Figures

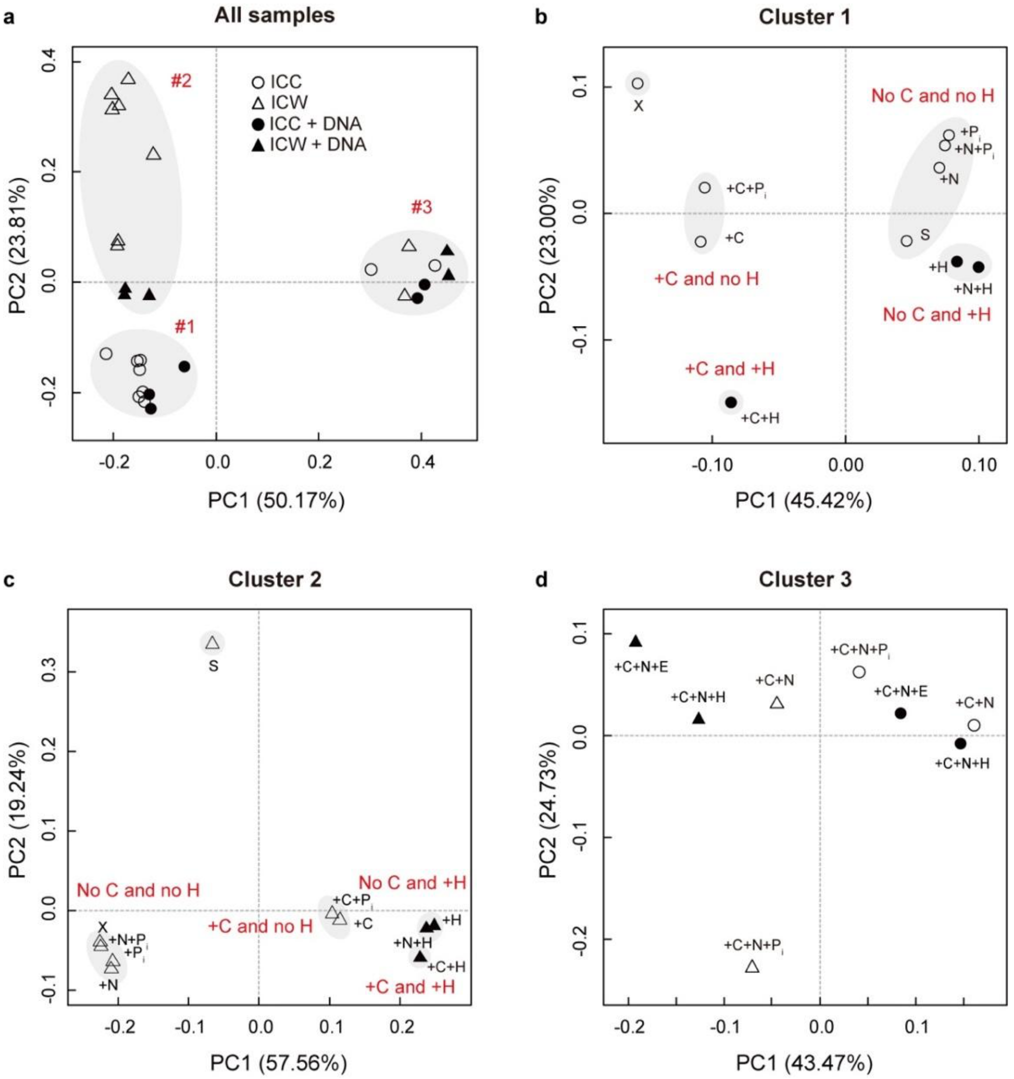


**Figure S1. Comparison of the 16S rRNA-based OTU compositions of the analyzed samples. (a)** Principal Coordinate Analysis of all samples using the pairwise Bray-Curtis dissimilarity as a distance metric. The distances were calculated using all OTUs in a sample. Circles denote DNA from cells representing the microbial community samples, while triangles refer to the dissolved eDNA in the water column samples. Within these, samples for treatments with added DNA are shown using filled symbols, while all remaining samples are shown using open symbols. "#1", "#2" and "#3" refer to Cluster 1, 2, and 3 discussed in the text. (**b**, **c** and **d**) Principal Coordinate Analysis of the samples within each of the three clusters, also using the pairwise Bray-Curtis dissimilarity as a distance metric. The panels use the same symbol notations as in panel **a**. Abbreviations for individual samples: X, pre-incubation community; S, starvation; "+" added nutrients, followed by a nutrient symbol of carbon (C), nitrogen (N), phosphorus ($P_i$), *Hfx. volcanii* DS2 DNA (H) or *E. coli dam$^-$/dcm$^-$* DNA (E).

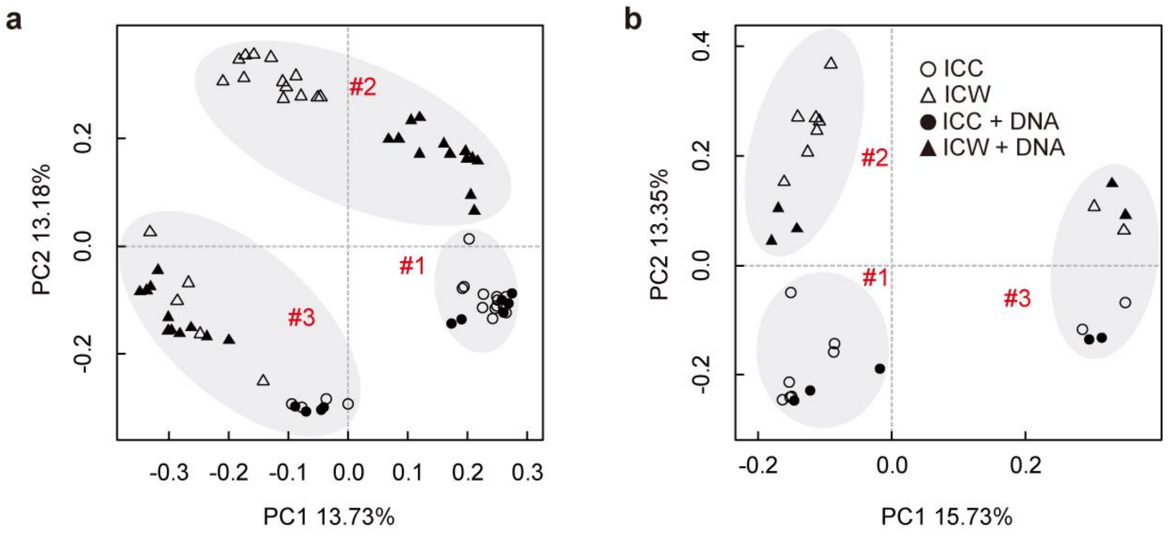


**Figure S2. Comparison of the OTU compositions of the analyzed samples taking into account only OTU richness.** Principal Coordinate Analysis of all samples using the Jaccard distances as a distance metric calculated for (**a**) *rpoB* and (**b**) 16S rRNA OTUs. Circles denote cells from the microbial community samples, while triangles refer to dissolved eDNA in the water column samples. Within these, samples for treatments with added DNA are shown using filled symbols, while all remaining samples are shown using open symbols. The "#1", "#2", and "#3" labels refer to the Cluster 1, Cluster 2 and Cluster 3 of samples, respectively, that are discussed in the text. Differences in OTU richness of the three clusters are significant (PERMANOVA: R = 0.65, P = 0.001 [rpoB] and R = 0.68, P = 0.001 [16S rRNA]).

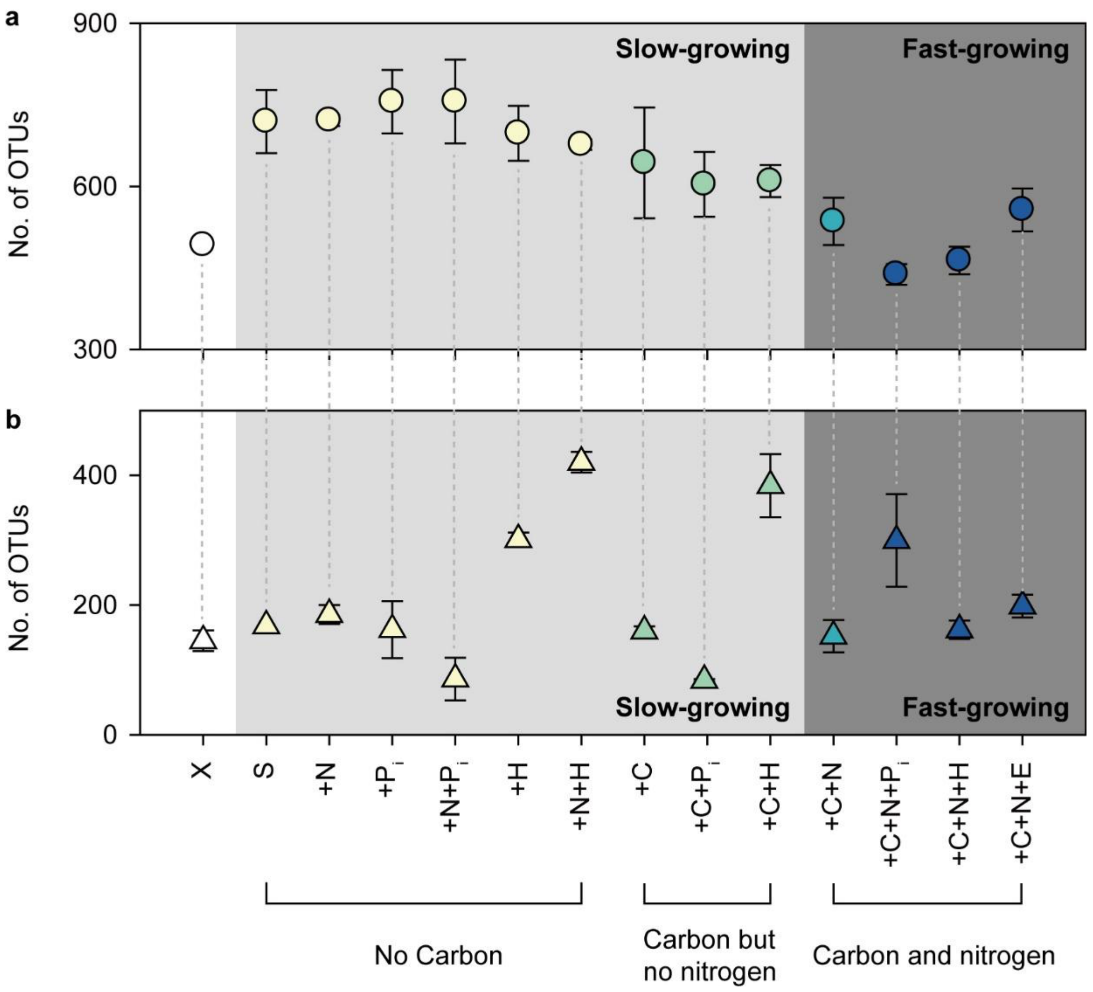


**Figure S3. Number of OTUs in microbial communities (panel a) and in eDNA from water columns (panel b).** For each sample, the average OTU number across replicates and standard error are displayed. Samples on X axis are listed in the same order as in **Fig. 3e**. The samples are further grouped based on whether the microcosms were supplemented with carbon and/or nitrogen sources.

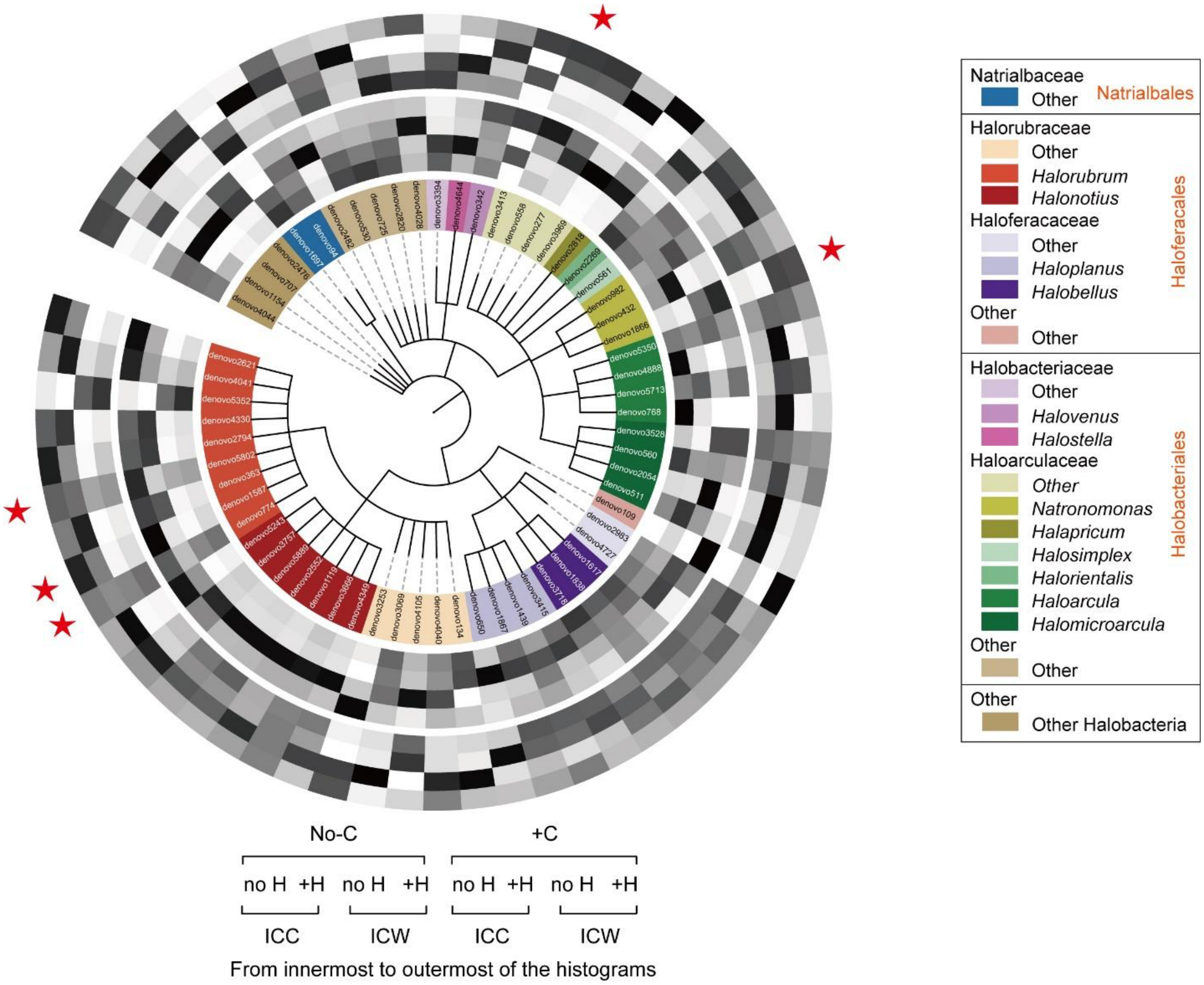


**Figure S4. Changes in ahOTU abundance in slow-growing communities and corresponding eDNA pool in response to addition of *Hfx. volcanii* DNA.** The ahOTUs are arranged into a cladogram according to the NCBI taxonomy, with relationships among ahOTUs from the same genus shown as unresolved. Each ahOTU is associated with a heatmap that summarizes its relative abundance across samples. The samples are arranged into two rings: those that came from microcosms with no notable growth (no C provided; inner ring) and with some growth (C provided; outer ring). Within each ring, the samples are further subdivided based on whether they come from cells (ICC) or dissolved in the water column (ICW) and if *Hfx. volcanii* DNA was provided (no-H/+H). Within each heatmap ring, the relative abundance of an ahOTU is normalized across four samples (see **Methods** for details) and visualized from white (low abundance) to black (high abundance). Stars denote examples of ahOTUs, whose eDNA is depleted in the water column when *Hfx. volcanii* DS2 DNA is not provided. The raw and normalized relative abundance values are provided in **Supplementary Table S7**.

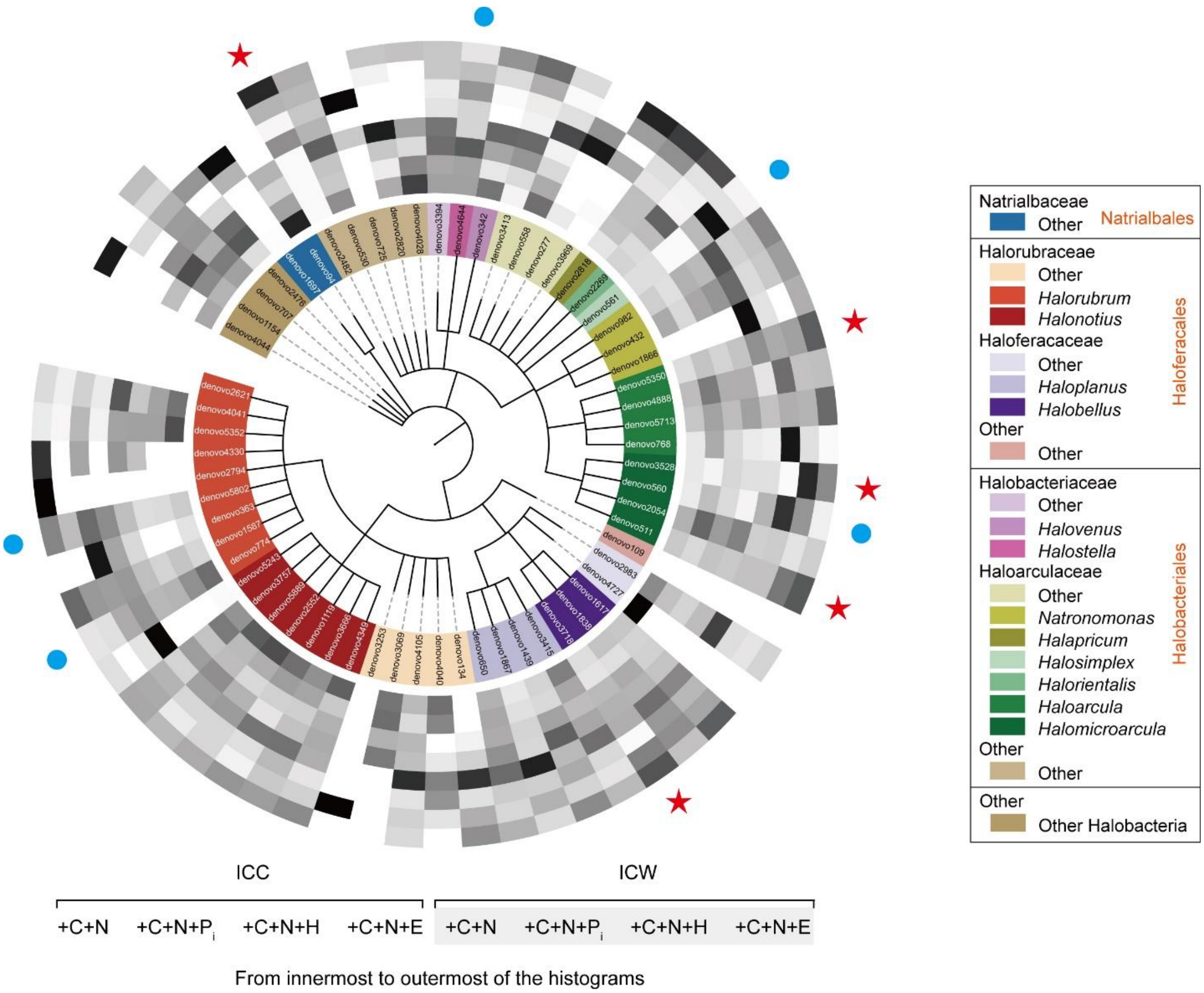


**Figure S5. Changes in ahOTU abundance in fast-growing communities and corresponding eDNA pool in response to provided phosphorus source.** The ahOTUs are arranged into a cladogram according to the NCBI taxonomy, with relationships among ahOTUs from the same genus shown as unresolved. Each ahOTU is associated with a heatmap that summarizes its relative abundance across samples. The relative abundance of an ahOTU is normalized across eight samples (see **Methods** for details) and visualized from white (low abundance) to black (high abundance). Red stars denote examples of ahOTUs whose eDNA is depleted in the water column when $P_i$ is not provided. Blue circles denote examples of ahOTUs whose eDNA is depleted in the water column even when $P_i$ is provided. The raw and normalized relative abundance values are provided in **Supplementary Table S9**.

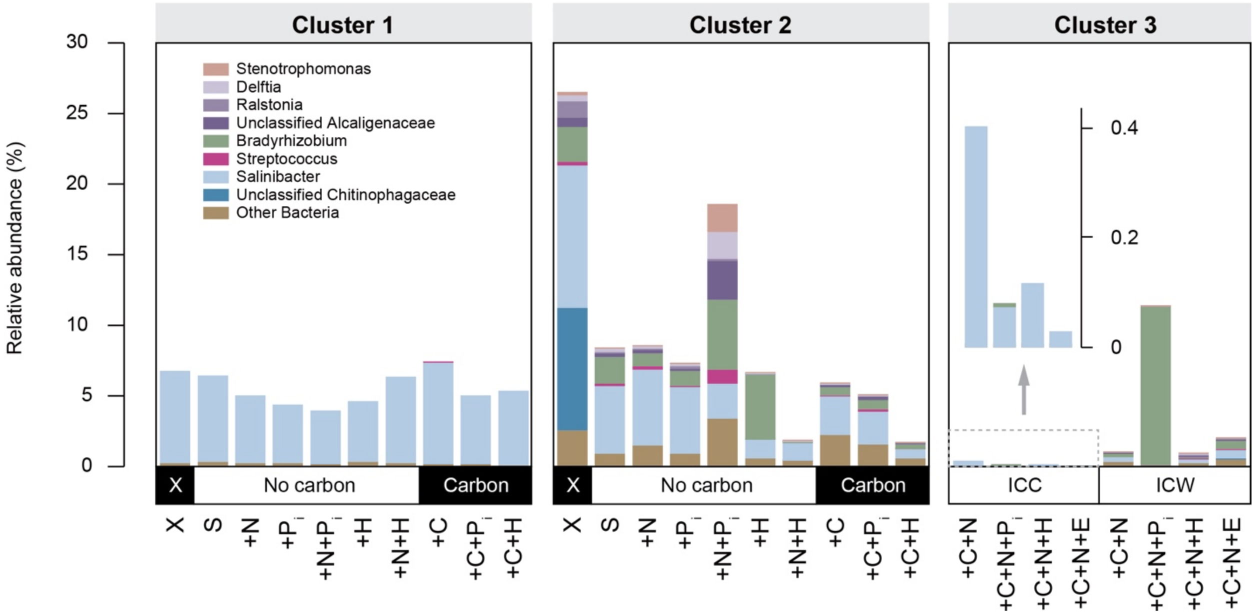


**Figure S6**. **Relative abundance of bacterial taxa across samples.** The samples are arranged according to the three clusters defined in **Figure 3.** For each sample, the height of a bar represents relative abundance of all bacterial OTUs in a sample. Within each bar, the relative abundance of shown genera or families is calculated as a sum of relative abundances of OTUs that constitute ≥5% of the bacterial fraction in at least one of 69 analyzed samples. The bacterial OTUs below this abundance cutoff are aggregated into "Other Bacteria" category. For sample notations shown on X-axes see **Figure S1** legend. Due to the overall low abundance of bacteria in the microbial community samples of Cluster 3, their relative abundances are additionally shown on a different scale as an inset.

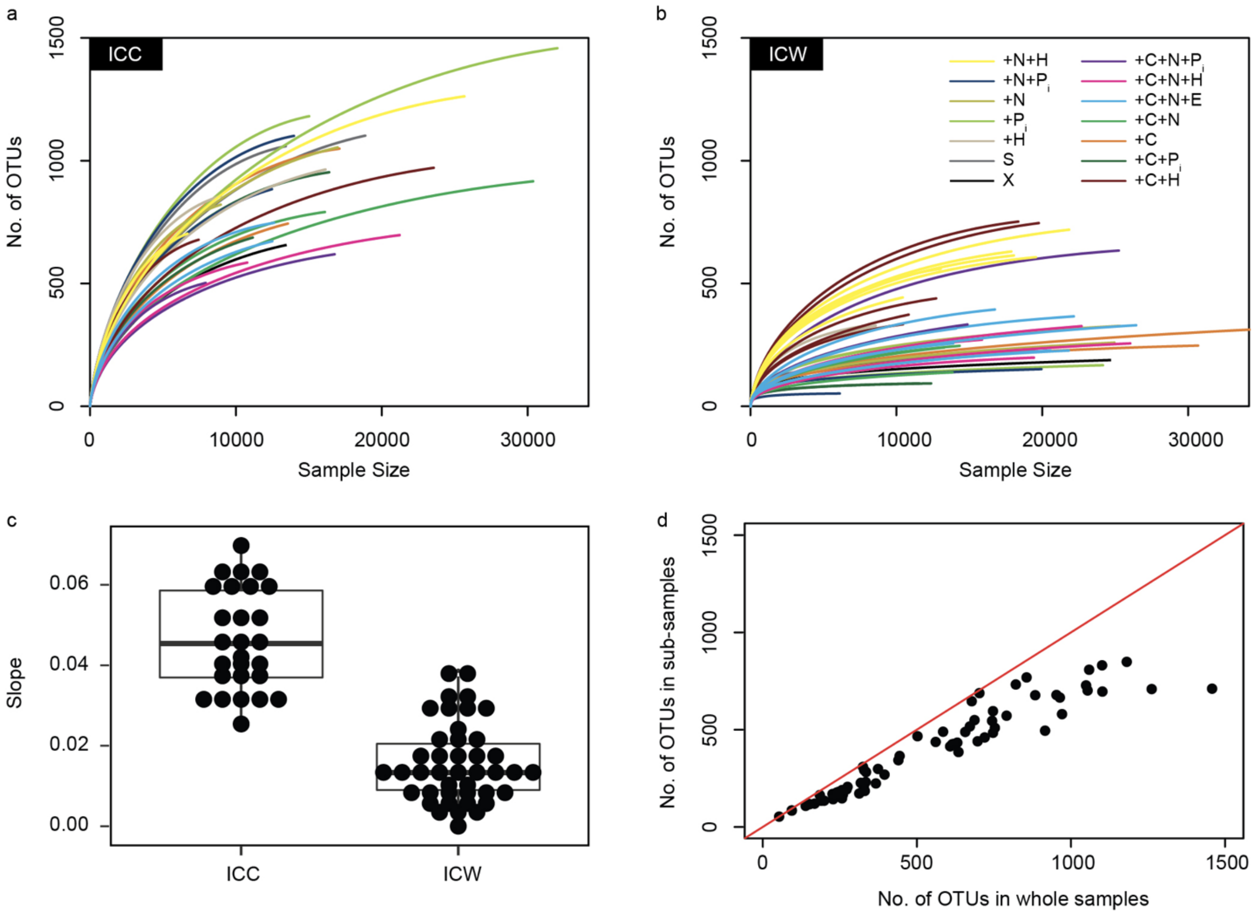


**Figure S7. Assessment of sequencing depth adequacy for *rpoB* data. Panels a** and **b**: Rarefaction curves for samples of microbial communities (ICC) and water columns (ICW). Sample size refers to the number of sequences after the quality control and merging of pairs. **Panel c**. Slopes of rarefaction curves calculated for the subsamples of 6 000 sequences that were randomly selected from each original sample. The box plot shows the upper quantile, median, and lower quantile of the data. **Panel d**. The relationship between the number of OTUs observed in whole samples and in the sub-samples of 6 000 sequences. Red line demarcates 1:1 relationship.

### Supplementary Results

##### Evaluation of sequencing depth adequacy

To evaluate if the amount of sequencing for each sample was sufficient to capture the diversity in microbial communities and water columns, we carried out a rarefaction analysis for the *rpoB*-based sequencing datasets. For the whole samples, rarefaction curves are plateauing (**Supplementary Figure S7a** and **S7b)**. Furthermore, for all sub-samples of 6 000 randomly selected sequences the slopes of rarefaction curves are below 0.07 (**Supplementary Figure S7c**), indicating that selected sub-samples reached saturation. Comparison of the number of OTUs in whole samples and the sub-samples of 6 000 sequences shows that selected subsamples in general well represent whole samples (**Supplementary Figure S7d**). Taken together, we conclude that the selected sub-samples, which were chosen to avoid biases in inter-sample comparison, adequately represent DNA content of the microcosms.

##### Additional details on the taxonomic composition of the microbial communities and eDNA pool varies across treatments

We note that almost all post-treatment communities have higher overall alpha-diversity than the pre-incubation community (**Supplementary Table S3**). Since post-treatment communities have descended from the pre-incubation community, these observations suggest that some OTUs had extremely low initial abundance in the pre-incubation community and therefore were not detected. An increase in abundance of these OTUs after treatments elevated them above detection limit.

Some DNA comes from OTUs that were either not observed in any of the ICC samples or were found in the ICC samples in low abundances. For example, DNA from two OTUs from bacterial families Alcaligenaceae and Chitinophagaceae is found only in the ICW samples (**Supplementary Table S4**), and therefore may represent eDNA that originated from living microorganisms elsewhere and was transported to the saltern environment. An OTU from the bacterial genus *Bradyrhizobium* and OTUs from the archaeal subphylum Nanohaloarchaeota may be recalcitrant-to-consumption eDNA that has slowly accumulated from the rare members in the pre-incubation community. *Bradyrhizobium* eDNA is abundant in three water column samples (**Fig. 2a**), but its cellular DNA is observed in only one ICC sample and at low abundance ("+C+N+$P_i$" treatment; **Supplementary Table S4**). Nanohaloarchaeota eDNA is present at >1% abundance in multiple water column samples, with the highest overall abundance (14%) found in the pre-incubation water column sample (ICW sample "X"; **Fig. 2a**), and although their cellular DNA is found in in all ICC samples as well, it is always present there at low abundances.

##### On differences between 16S rRNA- and *rpoB*-based taxonomic assignments

A few discrepancies were observed between relative abundances of some haloarchaeal OTUs in 16S rRNA and *rpoB*-based taxonomic assignments. Below we detail specific notable differences and explain possible underlying causes.

First, OTUs assigned to *Halomicroarcula* genus have much lower relative abundance in the 16S rRNA-based taxonomic assignments. We conjecture this is due to inability to distinguish amplified fragments of 16S rRNA sequences of *Halomicroarcula* and *Haloarcula* genera. For example, while the full-length 16S rRNA genes of *Halomicroarcula limicola* strain YGHS32 (NR_133757.1) and *Haloarcula marismortui* strain CGMCC1.1784 (NR_116086.1) are 96.4% identical, for the amplified region, the two sequences have 97.6% sequence identity, which is above the used OTU assignment cutoff of 97%. Therefore, the amplified *rpoB* fragments from these two genera would be placed into one OTU and assigned one or the other genus. Notably, across samples from the fast-growing communities, in which there is a high abundance of the OTUs from these genera, the sum of the relative abundances of *Haloarcula* and *Halomicroarcula* is on average 38.7% and 37.5% in *rpoB* and 16S rRNA-based analyses, respectively. This roughly equal representation of the two genera combined further support the proposed cause of the discrepancy.

Second, OTUs assigned to *Haloquadratum, Halohasta* and *Halomicrobium* genera are abundant in the 16S rRNA-based taxonomic assignments, but are represented by only few (if any) OTUs in the *rpoB*-based taxonomic assignments. We conjecture that this is due to reduced ability of the designed *rpoB* primers to amplify *rpoB* genes from these genera due to nucleotide mismatches in the non-degenerate bases of the primers. For example, there are 4-5 mismatches between the reverse primer sequence and the corresponding region of the *rpoB* sequence in four examined *Haloquadratum* genomes (NC_017459.1, NC_008212.1, ARPX00000000.1 and ARPY00000000.1). The *rpoB* gene sequence from the *Halohasta litchfieldiae* strain tADL genome (CP024845.1) and the reverse primer have two mismatches. Finally, the *rpoB* gene sequence from the *Halomicrobium* genomes (JN120806.1, NC_008212.1, ARPX00000000.1 and ARPY00000000.1) has one mismatch in both the forward and reverse primers. Note that to avoid mistaking sequencing errors for true mismatches, we limited the above analyses to the genomes that were estimated by CheckM [1] to have completeness of ≥ 80% and contamination of < 5%.

Third, the only ahOTU assigned to *Halosimplex* genus was present in the *rpoB*-based taxonomic assignments (although at an average relative abundance of only 0.3% across all samples) but absent in the 16S rRNA-based taxonomic assignments. The representative *rpoB* sequence of this OTU has 94.8% nucleotide identity and 100% coverage to the *rpoB* sequence in the *Halosimplex pelagicum* strain R2 genome (KF434759.1) and therefore likely belongs to the *Halosimplex* genus. In 16S rRNA dataset, there is an ahOTU that has 95.7% identity and 100% coverage to two database sequences that are classified as *Halosimplex* and *Halapricum* genera. Because of this ambiguity, the OTU has been classified at a higher taxonomic rank (as Haloarculaceae), but is likely to correspond to the *Halosimplex* OTU detected in the *rpoB* dataset.

### Supplementary Methods

*Quantification of nutrients in Isla Cristina water samples*

The samples (see the **Materials and Methods** section in the main text for details) were analyzed at the University of Connecticut Center for Environmental Sciences and Engineering (CESE). The change in nutrient concentrations before and after experiments was assessed by calculating log-odds ratios of the nutrient concentrations before and after experiments (**Supplementary Tables S1** and **S2**).

*Strains and growth conditions for collecting DNA to be supplied to the microcosms*

The strains *Haloferax volcanii* DS2 [2]and *Escherichia coli* with deleted *dam* and *dcm* methyltransferase genes (*E. coli dam⁻/dcm⁻*) were used in this study. All *Haloferax volcanii* strains were grown in Hv-YPC or Hv-CA medium described in the Halohandbook [3]. When needed, the media were supplemented with uracil (50µg/mL) and 5-fluoroorotic acid (5-FOA; 50µg/mL). Solid media for growth on Petri dishes was made by adding 2% (w/v) agar to the media. All *Escherichia coli* strains were grown in lysogeny broth (LB), which was made by adding 5g NaCl, 5g tryptone, and 2.5g yeast extract to deionized water with a final volume of 500 mL and a pH of 7.0. When necessary, LB was supplemented with ampicillin (100µg/mL). Solid media for growth on Petri dishes was made by adding 1.5% (w/v) agar to the medium.

*Incubation of treatment microcosms*

The microcosms were incubated at 42°C while shaking (200 RPM), and the growth of each microcosm was monitored approximately every 24 hours by measuring the optical density at 600 nm ($OD_{600}$), with Isla Cristina water used as a control. Once the microcosms reached stationary phase, they were centrifuged at 9 000x g for 45 minutes to pellet the cells, and the water from each tube was filter-sterilized (0.22µm) into new tubes. The cell pellets and filtered water samples were stored at -20°C until needed for DNA extraction and further analyses.

*DNA purification,* rpoB *and 16S rRNA genes amplification, and DNA sequencing*

To extract DNA for sequencing, the cell pellets were resuspended in 5mL of 10mM Tris-HCl buffer (pH 8.0) and treated with proteinase K (50µg/mL). The tubes were incubated overnight at 37°C to degrade the proteins, after which the DNA was precipitated via ethanol precipitation. The DNA samples were then purified with phenol-chloroform and chloroform extractions as needed, followed by a final ethanol precipitation on the remaining DNA. The concentration of the DNA samples was quantified via Qubit dsDNA BR assay (Invitrogen), and the 260/280 ratios were measured via Nanodrop. To extract DNA from the filtered water samples for sequencing, small aliquots (~20µL) of each sample were diluted to a NaCl concentration of ~0.2M, after which ethanol precipitation was performed to extract the DNA. The samples were then quantified via Qubit dsDNA HS assay (Invitrogen), and the 260/280 ratios were measured via Nanodrop. The samples were purified further via Amicon Ultra 30K centrifugal filter units (Millipore) to concentrate the samples.

For amplification of the *rpoB* gene from the archaeal class *Halobacteria*, we created a sequence alignment of *rpoB* genes from representative Halobacteria and based on the sequence conservation designed the following primers to 344F, ATCGACGCCGARGARGARGARGA and 634R, CATCACGGCGACGRYGAAGTTCTG. Amplification was done using Phusion polymerase (Thermo Scientific) in the supplied HF buffer with the addition of 10% (v/v) DMSO. The PCR reaction was incubated at 98˚C for 30 s initial denaturation, then 35 cycles of 5 s at 98.0°C, 15 s at 50.5°C and 15 s at

72.0°C, followed by a final extension of 5 minutes at 72.0°.

For the 16S rDNA gene, the universal primers 515F/806R (V4 region, AGAGTTTGATCMTGGCTCAG/ ATTACCGCGGCTGCTGG, 0.8 picomole each 515F and 806R with Illumina adapters and 8 base pair dual indices [4]) were amplified in triplicate 15ul reactions using GoTaq (Promega) with the addition of 10µg BSA (New England BioLabs). To overcome initial primer binding inhibition due to the mismatch of the majority of the primers to the template priming site, 0.1 femtomole 515F and 806R without the barcodes and adapters was added. The PCR reaction was incubated at 95˚C for 2 minutes, the 30 cycles of 30 s at 95.0°C, 60 s at 50.0°C and 60 s at 72.0°C, followed by final extension as 72.0°C for 10 minutes. PCR products were pooled for quantification and visualization using the QIAxcel DNA Fast Analysis (Qiagen). PCR products were normalized based on the concentration of DNA from 350-420 bp then pooled using the QIAgility liquid handling robot. The pooled PCR products were cleaned using the Mag-Bind RxnPure Plus (Omega Bio-tek) according to the manufacturer's protocol.

The purified amplicon products were used to construct 69 *rpoB*-based libraries (1-2 libraries for the initial sample and 1-7 libraries for each microcosm treatment) and 28 16S-rRNA-based libraries (one library for the initial sample and each microcosm treatment). The libraries were sequenced on an Illumina MiSeq instrument in the PE 250 mode using v2 2x250 base pair kit at the Microbial Analysis, Resources and Services facility at the University of Connecticut (Storrs, CT).

*Sequence quality control and assembly*

From the raw reads, the bases with *phred* scores below 20 were trimmed using the in-house Perl script. The paired reads were merged using FLASH v1.2.11 [4] with "-m 10 -x 0.25" options. To eliminate poor quality assemblies, the bases with average *phred* scores below 20 in a sliding window of 50 bp were trimmed using the "*trim.seqs*" script of Mothur v1.39.5 [5]. Both the forward and reverse primers were removed from the merged pairs. Sequences that were found only once across all samples were removed.

*Grouping of merged pairs into the operational taxonomic units and assignment of taxonomic affiliations*

The assembled *rpoB*- and 16S rRNA-based sequences were clustered into operational taxonomic units (OTUs) by applying *uclust* v1.2.22q algorithm [6] at the 95% and 97% nucleotide identity level, respectively, as implemented in the "*pick_otus.py*" script of QIIME v1.9.1 [7]. The sequence of an OTU designated by QIIME during the clustering step as 'the cluster seed' was selected as a representative sequence of the OTU. Taxonomy of the representative sequence was assigned using the Ribosomal Database Project classifier v2.2 [8] with the minimum confidence set at the 80% threshold. This taxonomic assignment was considered to be taxonomy of the whole OTU. The *rpoB* sequences that could not be assigned to *Halobacteria* were removed from further analyses. For the 16S rRNA classifications, RDP release 11, update 5 served as the reference database (downloaded on Aug 27$^{th}$, 2020) [9]. For the *rpoB* classifications, a database of 627 sequences was constructed, which consists of the *rpoB* genes from 346 complete and draft halobacterial genomes available in GenBank on June 13$^{th}$, 2019 and the *rpoB* genes from the *nt* database (downloaded on November 16$^{th}$, 2017) that matched the 344F/634R primers. The database is available in **Supplementary Data**.

*Evaluation of sequence depth of rpoB-based sequencing*

Rarefaction analysis for each assembled sample of *rpoB*-based sequences was carried out using *vegan* v2.5-6 package in R v3.5.3. The rarefaction curves were calculated using the "*rarecurve*" function. The slopes of the rarefaction curves were calculated using "*rareslope*" function. The expected OTU richness was calculated using the "*rarefy*" function.

*Heterogeneity of 16S rRNA gene copies in the genomes of haloarchaea*

Four hundred eighty-nine available complete and draft genomes from both class *Halobacteria* and phylum *Nanohaloarchaeota* were downloaded from the NCBI Refseq database on May 21$^{th}$, 2019. From each genome, all copies of the 16S rRNA genes were extracted using the "*ssu_finder*" program of CheckM v1.0.13 [1]. Sequences with length < 1 000 bp were removed from further analyses. This selection procedure resulted in 478 16S rRNA gene sequences from 346 genomes, with one to six copies of the gene per genome. For the genomes with multiple copies of the 16S rRNA gene, the percent identities among copies and the alignment coverage was extracted from the aligned regions of the pairwise local alignments reconstructed using BLASTN v2.6.0+ [10].

*Quantification of similarities and differences in taxonomic composition across and within samples*

Relative abundance of an OTU in a sample was calculated as the number of sequences in the OTU divided by the total number of sequences in the sample. Overall taxonomic composition of each sample, which are shown on **Fig. 2**, **3e** and **S6,** was calculated using all sequences in each sample, averaging across treatment replicates when available. However, to avoid the possible biases due the unevenness of sample sizes during inter-sample comparison, for the comparisons described in the remainder of this section, each sample was randomly sub-sampled to the 6 000 sequences for *rpoB* and to the 15 000 sequences for 16S rRNA gene datasets using "single_rarefaction.py" script of QIIME v1.9.1 [7].

The OTU diversity within a sample was summarized by calculating Shannon [11, 12], Simpson [13, 14], Observed species, Chao1 [15] indices of α-diversity. The OTU diversity among samples was summarized by calculating Bray-Curtis [16] and Jaccard [17] dissimilarity indices of β-diversity. All indices were calculated using the "alpha_diversity.py" and "beta_diversity.py" scripts of QIIME v1.9.1 [7].

Grouping of samples into clusters was performed via Principal Coordinates Analysis (PCoA), using Bray-Curtis and Jaccard dissimilarity matrices as inputs, as implemented in the *vegan* package v2.3-5 in R v3.2.5. To test the difference among clusters identified in the PCoA analysis, permutational multivariate analysis of variance (PERMANOVA, as implemented in the "*adonis*" function in R) was performed on the Bray-Curtis distance matrix, using 999 permutations for each analysis. Significance of the differences of OTU relative abundances among different groups of samples and among various treatment combinations was tested using the nonparametric Mann-Whitney *U* test, with confidence intervals set at 99% significance. To avoid inflation of false positives due to the multiple testing, P values were corrected using Benjamini-Hochberg correction ($P < 0.05$), as implemented in the "*p.adjust*" program in R.

Relative abundance of ahOTUs across Clusters 1, 2 and 3 shown on **Fig. 4** was visualized in ternary plots using OriginPro 2017 v. b9.4.0.220 (OriginLab Corporation).

*Examination of the impact of provided nutrients on community composition using linear regression*

Linear regression modeling was conducted with provided nutrients (C, N, $P_i$ and $P_o$) as explanatory variables and number of OTUs in each sample as dependent variables. For each sample, the nutrients provided to the microcosm were recoded as follows: addition of C or N was treated as binary variable, while addition of P was recoded into a contrast matrix, due to the three possible values (no provided source of P, provided $P_i$ and provided $P_O$). The regression analysis was carried out using the "*lm*" function in *stats* R package v3.2.5.

*Visualization of changes in relative abundances of the ahOTUs in response to provided phosphorus*

Changes in ahOTU abundance in microcosms and eDNA pools in response to addition of DNA and inorganic phosphorus were visualized as heatmaps mapped to a cladogram of relationships among ahOTUs (**Supplementary Fig. S4** and **S5**). The cladogram of ahOTU relationships was generated using NCBI Taxonomy hierarchy based on assigned taxonomic information of ahOTU obtained as described in the "*Grouping of merged pairs into the operational taxonomic units and assignment of taxonomic affiliations*" section. Due to large differences in relative abundances of ahOTUs across examined samples, relative abundances of each ahOTU across a specified group of samples were normalized using the "*scale*" function in *base* R package v3.2.5.

*Prediction of provided nutrients from taxonomic composition of a sample*

The taxonomic composition of a sample was defined as the relative abundance of ahOTUs in the original (i.e., not sub-sampled) samples. All samples were partitioned into five categories based on combination of nutrients provided in the experiments: "No C was provided" (no C, 30 samples), "C but no N were provided" (+C, 15 samples), "Only C and N were provided" (No P, 5 samples), "C, N and DNA were provided" (+H/+E, 15 samples), and "C, N and $P_i$ were provided" (+$P_i$, 4 samples).

Sixty-nine samples were randomly divided into training and validation datasets, with 56 and 13 samples, respectively. Due to the variable number of samples in each of the five categories, 56 samples were required to represent each of the five categories by at least one sample. This training and validation dataset selection procedure was repeated 1 000 times. For each of the 1 000 training/validation datasets, random forest classifier [18] was built and trained using 20-fold cross-validation. The average prediction accuracy of the trained classifier was computed as the mean of the 1 000 repeats. Additionally, out-of-bag (OOB) error estimate was used to assess the internal generalization error [19]. All analyses were carried out using the *randomForest* package v. 4.6.14 in R version 3.5.1.

The mean decrease in accuracy for each ahOTU was extracted from the classifier output, and 24 ahOTUs that contribute ≥1% of the prediction accuracy were selected. The statistical significance of the change in the relative abundances of the 24 ahOTUs among the five categories was assessed using one-way ANOVA test, as implemented in the "*aov*" function in *stats* R package v3.2.5. To illustrate the differential presence of the 24 ahOTUs across five categories, the relative abundance values were transformed into color categories and visualized using the "*heatmap.2*" function in the *gplots* R package v. 3.0.1.

The performance of classifier trained on the real data was compared to that of a classifier trained on the re-shuffled samples, which were obtained by random re-assignment of the treatment labels among samples with taxonomic composition of each sample kept intact. The significance of classifier's accuracy for real and re-shuffled datasets was assessed using Student's *t* test (testing if the means of two sets are significant different) and Kolmogorov-Smirnov test (testing the probability distribution pattern between two samples), as implemented in the "*t.test*" and "*ks.test*" functions in *stats* R package v3.2.5.

To examine the sensitivity of the classifier's predictive capacity to the size of the training dataset, the training and cross-validation of the classifier was repeated using different proportions of samples. Specifically, between 50 and 95% of the 69 samples, examined at 5% increments and rounding down to the closest integer value to obtain the integer number of samples, were selected as training datasets. The remaining samples were used for cross-validation. Additionally, the "leave-one-out" procedure [20] was used. For each dataset size, the sample selection procedure was repeated 100 times and also required training dataset to contain at least one sample for each of the five categories. The cross validation and accuracy calculations were performed as described above.
